# Age-determined expression of priming protease TMPRSS2 and localization of SARS-CoV-2 infection in the lung epithelium

**DOI:** 10.1101/2020.05.22.111187

**Authors:** Bryce A. Schuler, A. Christian Habermann, Erin J. Plosa, Chase J. Taylor, Christopher Jetter, Meghan E. Kapp, John T. Benjamin, Peter Gulleman, David S. Nichols, Lior Z. Braunstein, Alice Hackett, Michael Koval, Susan H. Guttentag, Timothy S. Blackwell, Vanderbilt COVID-19 Consortium Cohort, Steven A. Webber, Nicholas E. Banovich, Jonathan A. Kropski, Jennifer M. S. Sucre, HCA Lung Biological Network

**Affiliations:** Department of Pediatrics, Vanderbilt University Medical Center, Nashville, TN; Division of Allergy, Pulmonary and Critical Care Medicine, Department of Medicine, Vanderbilt University Medical Center, Nashville, TN; Division of Neonatology, Department of Pediatrics, Vanderbilt University Medical Center, Nashville, TN; Department Pathology, Microbiology and Immunology, Vanderbilt University Medical Center, Nashville, TN; Department of Radiation Oncology, Memorial Sloan Kettering Cancer Center, New York, New York; Division of Pulmonary, Allergy, Critical Care, and Sleep Medicine, Emory University, Atlanta, GA; Department of Cell Biology, Emory University, Atlanta, GA; Department of Cell and Developmental Biology, Vanderbilt University, Nashville, TN; Department of Veterans Affairs Medical Center, Nashville, TN; Department of Medicine, Vanderbilt University Medical Center, Nashville, TN; Translational Genomics Research Institute, Phoenix, AZ

## Abstract

The SARS-CoV-2 novel coronavirus global pandemic (COVID-19) has led to millions of cases and hundreds of thousands of deaths around the globe. While the elderly appear at high risk for severe disease, hospitalizations and deaths due to SARS-CoV-2 among children have been relatively rare. Integrating single-cell RNA sequencing (scRNA-seq) of the developing mouse lung with temporally-resolved RNA-in-situ hybridization (ISH) in mouse and human lung tissue, we found that expression of SARS-CoV-2 Spike protein primer *TMPRSS2* was highest in ciliated cells and type I alveolar epithelial cells (AT1), and *TMPRSS2* expression was increased with aging in mice and humans. Analysis of autopsy tissue from fatal COVID-19 cases revealed SARS-CoV-2 RNA was detected most frequently in ciliated and secretory cells in the airway epithelium and AT1 cells in the peripheral lung. SARS-CoV-2 RNA was highly colocalized in cells expressing *TMPRSS2.* Together, these data demonstrate the cellular spectrum infected by SARS-CoV-2 in the lung epithelium, and suggest that developmental regulation of *TMPRSS2* may underlie the relative protection of infants and children from severe respiratory illness.

## Introduction

The emergence of the SARS-CoV-2 novel coronavirus has led to a global pandemic (COVID-19), with more than 18 million cases as of August 2020(1). While the global burden of disease has overwhelmed healthcare systems around the world, available data suggest that severe illness and death from COVID-19 are rare in the pediatric population(2). One aspect of the COVID-19 pandemic that has eluded explanation is the striking diversity of clinical phenotypes accompanying SARS-CoV-2 infection, ranging from asymptomatic carriage to life threatening multi-organ failure(3–8). Morbidity and mortality appear most severe among the elderly(9, 10), while infection rates and hospitalizations among infants and children are substantially lower(1). In a recently published study from China, 90% of children infected with SARS-CoV-2 exhibited mild symptoms or were asymptomatic(2), and neonatal infections are limited to a handful of case reports. This led us to hypothesize that host factors determining SARS-CoV-2 cellular infectivity and viral attachment in the respiratory epithelium may be developmentally regulated. Differential expression of these viral attachment factors across lung development may provide biologic rationale for the variability in COVID-19 presentation.

## Results and Discussion

To investigate how the expression of genes associated with SARS-CoV-2 susceptibility change during lung development, we analyzed a previously unpublished scRNA-seq dataset profiling the epithelial and stromal cells in the developing mouse lung at five timepoints ranging from embryonic day 18 (E18) to postnatal day 64 (P64) (Fig. 1A-B, Fig. S1A-B). We interrogated expression profiles of genes linked to SARS-CoV-2 infectivity by analyzing a total of 67,629 cells across these 5 timepoints (Fig. 1C-D). Previous work has suggested that SARS-CoV-2 gains cellular entry by binding ACE2 on the cell surface(11–13), then the spike-protein undergoes a protease-mediated cleavage facilitating fusion with the cell membrane (14). TMPRSS2 is the canonical protease mediating cellular entry for coronaviruses including SARS-CoV-2(14), although it should be noted that there are reports the SARS-CoV-2 spike protein may be cleaved by other proteases(13). Consistent with recent reports analyzing single-cell transcriptomic data of the lung and other organs (15–19), we observed that during lung development, expression of *Ace2* was generally low, was largely restricted to epithelial cells, was most frequently and most highly expressed in secretory cells (Fig. 1E), and was expressed in a small subset of alveolar type 2 (AT2) cells. In contrast, *Tmprss2* was expressed broadly in the epithelium and was most highly expressed in ciliated cells and alveolar type 1 (AT1) cells (Fig. 1E). A small proportion of fibroblasts and pericytes expressed *Ace2*, but there was minimal *Ace2* or *Tmprss2* in endothelial or other stromal cells (Fig. 1E). Examining the relative expression of *Tmprss2* and other putative priming proteases across developmental time (including furin and cathepsin B), we observed that specifically in ciliated airway epithelial cells, *Tmprss2* expression was significantly higher at P64 compared to all earlier developmental timepoints; a similar pattern was observed for *Ctsb*. In AT1 cells, *Tmprss2* expression generally increased during alveolarization into adulthood.

**Figure 1.**
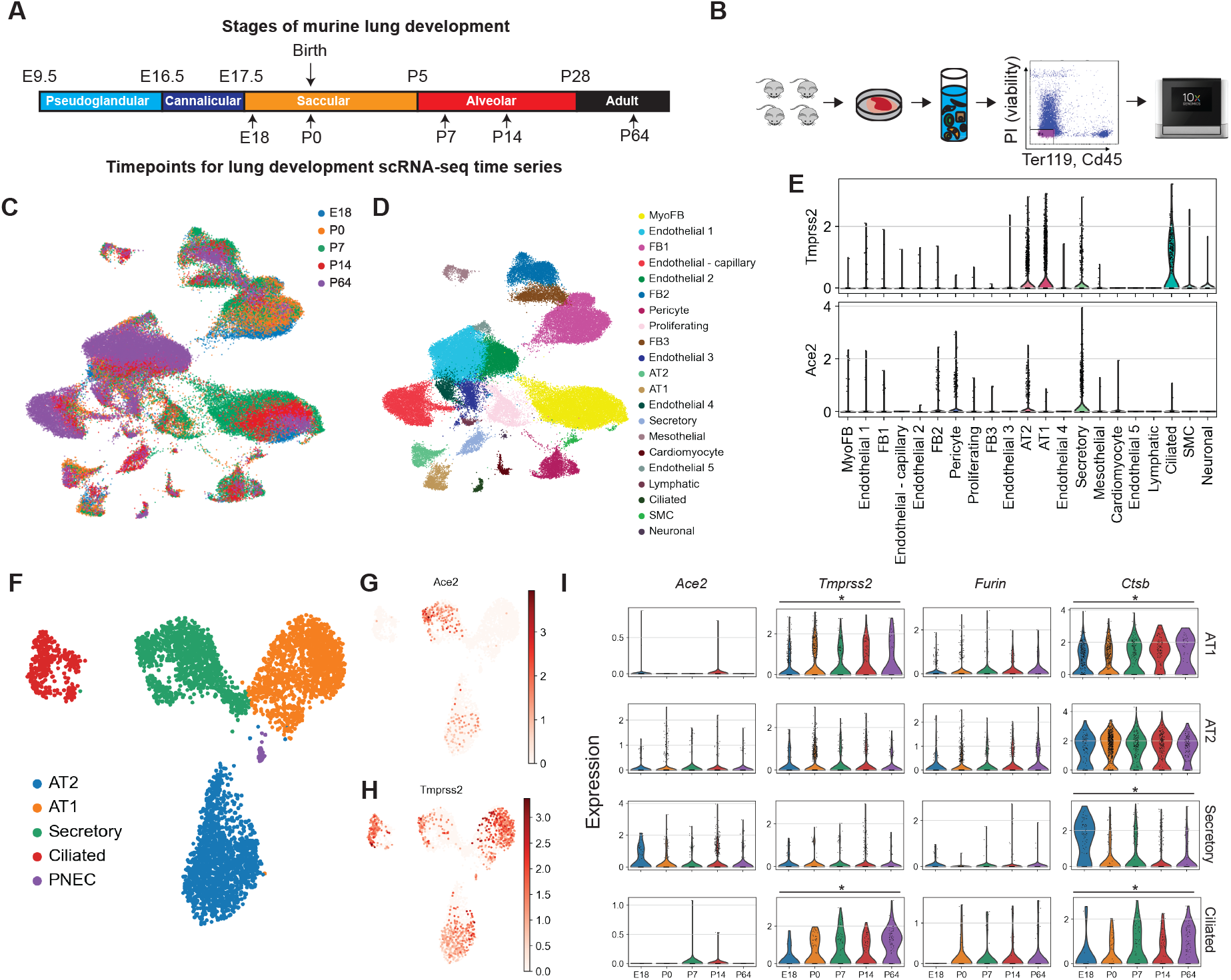
Time-series scRNA-seq of the developing mouse lung. A) Overview of mouse lung developmental stages and timepoints sampled for scRNA-seq time-series. B) Workflow of scRNA-seq time series. Single cell suspensions were generated from at least 4 mice at each timepoint. Viable, Cd45-, Ter119-cells were selected and underwent scRNA-seq library preparation using the 10X Genomics Chromium 5’ platform. c-d) UMAP embedding of 67,629 cells annotated by C) developmental timepoint and D) cell-type. E) Violin plot depicting expression of key SARS-CoV2 receptor *Ace2* and coreceptor *Tmprss2* across cell types in the jointly analyzed dataset. F) UMAP embedding of 4,369 epithelial cells after subsetting and reclustering. G,H) UMAP plots depicting relative expression of g) *Ace2* and h) *Tmprss2*. i) Relative expression of putative SARS-CoV-2 priming proteases across developmental time. \**p*<3.125×10^−3^ by ANOVA across developmental timepoints.

To provide tissue validation and to spatially and temporally localize expression of *Tmprss2,* we performed RNA-ISH using formalin fixed paraffin-embedded (FFPE) lung tissue from the same developmental time points used for scRNAseq, as well as lung tissue from 1yr and 2yr old mice. Colocalizing *Tmprss2* with *Scgb1a1* (secretory cells*), Foxj1* (ciliated cells), *Sftpc* (AT2 cells), and *Hopx* (AT1 cells) (Fig. 2A), we observed an age-dependent progressive increase in the proportion of *Tmprss2+* cells among all cell types (Fig. 2B). Further, in AT1 cells and ciliated cells, there was a marked increase in relative *Tmprss2* expression across developmental time that was most striking at 1 and 2 years of age (Fig. 2B). *Tmprss2* exhibited relatively low expression in secretory cells although levels increased in adult and aged mice. AT2 cells showed even lower expression of *Tmprss2* when compared with other epithelial cells, with little detectable increase in expression across lung development. Notably, there was minimal detection of *Tmprss2* prenatally at E18, and relatively low levels in the saccular stage at P0 in all epithelial cell types. Combining RNA ISH with immunofluorescence for TMPRSS2 protein showed concordance between expression of the *Tmprss2* gene and the TMPRSS2 protein (Fig. S2), and the localization of TMPRSS2 protein expression in ciliated cells expressing *Foxj1*was confirmed (Fig. S2). Consistent with our transcriptomic data, basal *Ace2* expression detected by RNA-ISH was low with little change during development and aging (Fig. S3).

**Figure 2.**
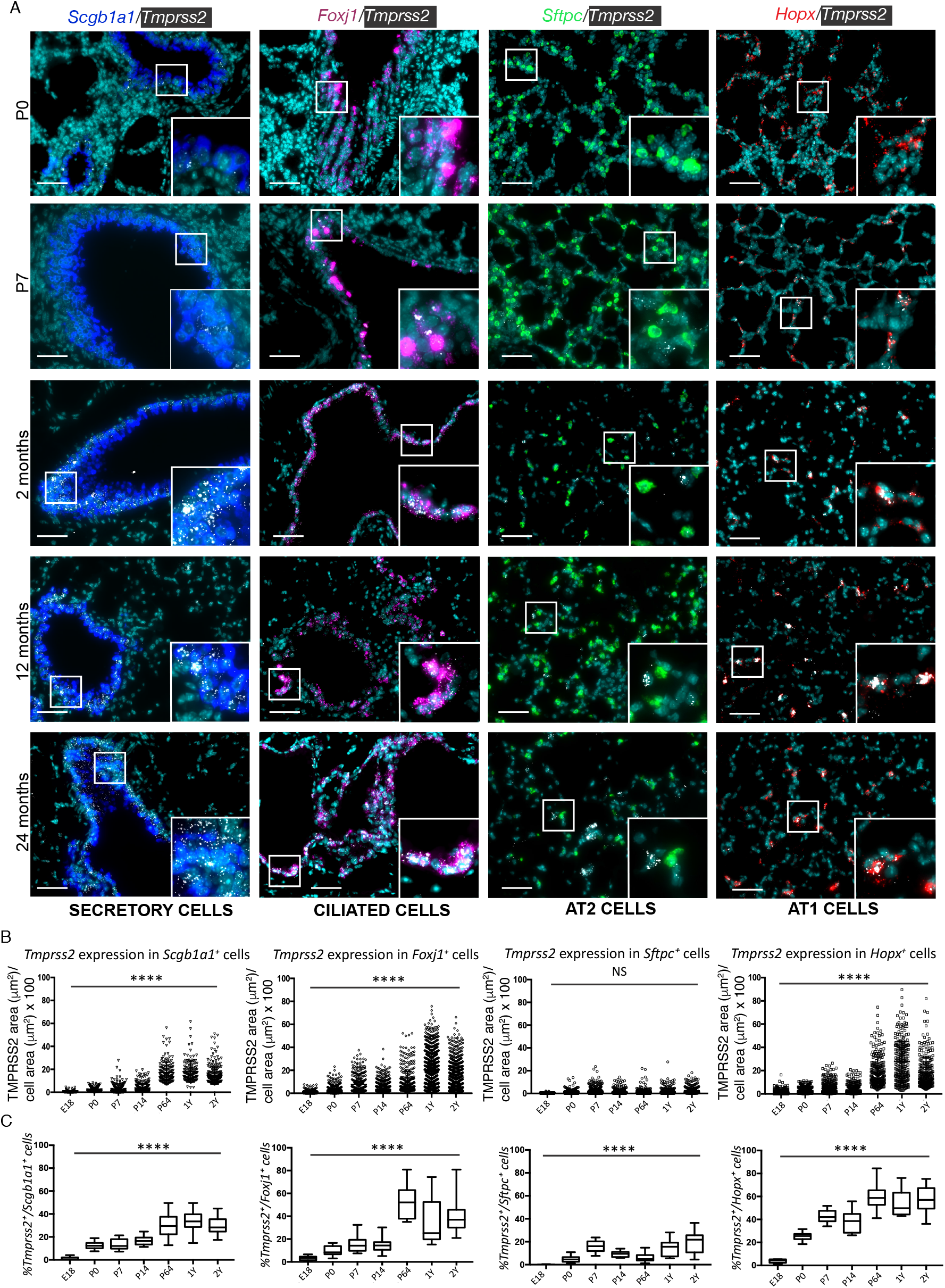
Spatial and temporal localization of *Tmprss2* expression across lung development. A) RNA in situ hybridization (ISH) of *Tmprss2* expression (white) with epithelial cell markers *Scgb1a1* (secretory cells, blue), *Foxj1* (ciliated cells, magenta), *Sftpc* (surfactant protein C, AT2 cells, green), *Hopx* (type 1 alveolar epithelial cells, red). Formalin fixed paraffin embedded tissue from lungs at timepoints E18, P0, P7, P14, P64, 12 months, and 24 months was used, with representative image data from timepoints P0, P7, P64, 12 months, and 24 months shown in the figure. Lungs from 3 mice at each time point were used, with ten 40X images obtained per slide. Scale bar = 100μm. B, C) Quantification of *Tmprss2* expression in each epithelial subtype across development measured as: B) a percentage of cellular area covered by *Tmprss2* probe for each cell expressing both *Tmprss2* and the epithelial cell marker; all data points are shown with mean +/− standard deviation (SD) and C) percentage of cells expressing the epithelial cell marker that also express *Tmprss2,* with positive *Tmprss2* expression defined has having 5 or more copies of *Tmprss2* probe; box and whisker plots are shown with mean and error bars reflecting minimum and maximum values for each timepoint. Greater than 1000 cells were counted at each timepoint. \*\*\*\**p*<0.0001 by one-way ANOVA.

To determine whether *TMPRSS2* is also developmentally regulated in humans, we examined *TMPRSS2* expression by RNA-ISH across the human lifespan. We defined infants as individuals up to 2 years of age (n=7), children between the ages of 3 and 17 (n=9). Adult specimens were from subjects aged 54-69 (never smokers, n=4). Infants expressed *TMPRSS2* at very low levels across all four epithelial lineages evaluated, while children exhibited similarly low levels of *TMPRSS2* in secretory and alveolar epithelial cells with a significant increase in *FOXJ1+* ciliated cells (Fig. 3A-B). Adult subjects had higher *TMPRSS2* expression in secretory, ciliated, and AT1 cells relative to both pediatric groups with very little *TMPRSS2* expression in AT2 cells (Fig. 3B). As with the murine experiments, combining RNA ISH with immunofluorescence for TMPRSS2 protein showed concordance between expression of the *Tmprss2* gene and the TMPRSS2 protein (Fig. S3). The localization of TMPRSS2 protein expression in ciliated cells expressing *Foxj1,* and the substantial increase of TMPRSS2 protein expression with aging was confirmed (Fig. S4). Human lung tissue demonstrated low levels of *ACE2* expression across in infants, children, and adult samples (Fig S5). These data are broadly consistent with results of a single-nucleus RNA-seq (snRNA-seq) study of infants, children and adults up to age 30 that was reported while this manuscript was being finalized, though notably that study only had 3 subjects per group and did not show tissue validation of their findings(20).

**Figure 3.**
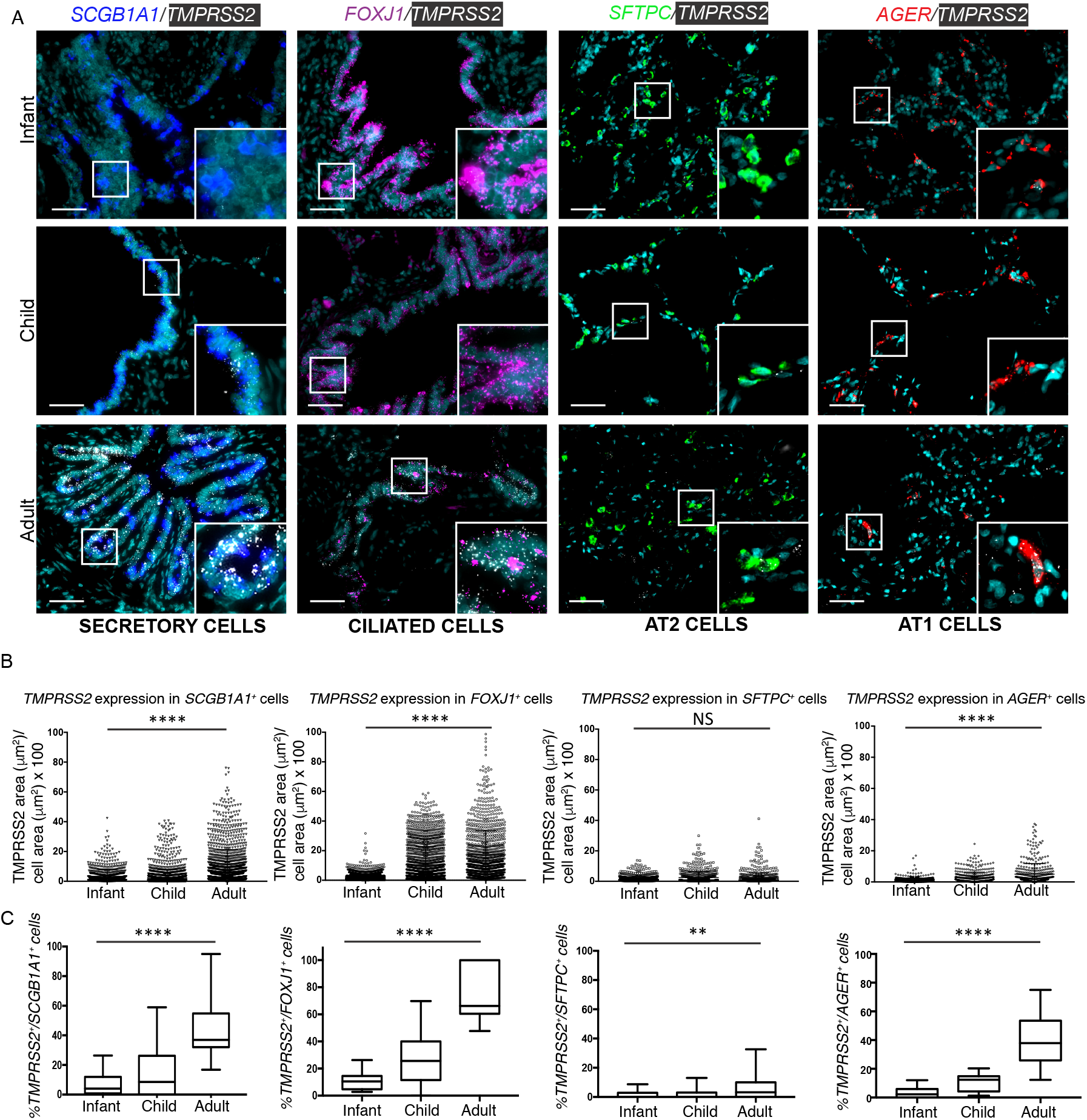
Spatial and temporal localization of *TMPRSS2* expression in the lung across the human lifespan. A) RNA *in-situ* hybridization (ISH) of *TMPRSS2* expression (white) with epithelial cell markers *SCGB1A1* (secretory cells, blue), *FOXJ1* (ciliated cells, magenta), *SFTPC* (surfactant protein C, AT2 cells, green), *AGER* (AT1 cells, red). Formalin fixed paraffin embedded tissue from 20 human lungs between the ages of birth and 69 years was analyzed. We defined infants as birth-2 years, children 3years-17 years, and adult specimens were between 53-69 years of age. Ten 40X images were obtained per slide for analysis. Scale bar = 100 μm. B, C) Quantification of *TMPRSS2* expression in each epithelial subtype across development as measured as: B) a percentage of cellular area covered by *TMPRSS2* probe for each cell expressing both *TMPRSS2* and the epithelial cell marker and C) percentage of cells expressing the epithelial cell marker that also express *TMPRSS2,* with positive *TMPRSS2* expression defined has having 5 or more copies of *TMPRSS2* probe. Greater than 1000 cells were counted at each timepoint. \*\*\*\**p*<0.0001, \*\**p*<0.01 by one-way ANOVA.

To understand the relevance of our findings in the setting of COVID-19 infection, we analyzed autopsy specimens from the lungs of patients who died from complications of COVID-19 (n=3). Using RNA-ISH, we identified the presence of SARS-CoV-2 RNA in both the large airway and lung parenchyma. SARS-CoV-2 was localized to epithelial cells expressing *SCGB1A1* (secretory)*, FOXJ1* (ciliated), and *AGER* (AT1). Surprisingly, despite analyzing > 150 *SFTPC* positive cells (corresponding to AT2), none contained detectable SARS-CoV-2 RNA by this assay (Fig 4A-B). In large airways (mainstem bronchus), SARS-CoV-2 co-localized with *TMPRSS2 in SCGB1A1+* secretory cells as well as *SCGB1A1*-negative (presumably ciliated) cells (Fig. 4C), and SARS-CoV-2 RNA was highly colocalized in cells expressing *TMPRSS2* (Fig. 4D). Since autopsy tissue likely represents end-stage disease, the surprising absence of SARS-CoV-2 in AT2 cells could be explained by changes in viral replication during the course of the illness and/or increased cytotoxicity of infected AT2 cells; alternatively direct viral infection of AT2 cells may not play as significant role in the disease pathophysiology as has been hypothesized. It is also possible that SARS-CoV-2 is localized to other cells in the lung that we did not look for, including inflammatory cells. These data do indicate that secretory, ciliated, and AT1 cells had the highest levels of *TMPRSS2* expression and that these cell types all harbored SARS-CoV-2 RNA in lung tissue from severely infected patients, similar to data observed in an *ex-vivo* infection model(21). Immunofluorescence for TMPRSS2 protein showed localization of SARS-CoV-2 virus in cells expressing TMPRSS2. Together, these data suggest that priming protease TMPRSS2 expression may be a crucial determinant of SARS-CoV-2 infectivity in the lower respiratory tract.

**Figure 4.**
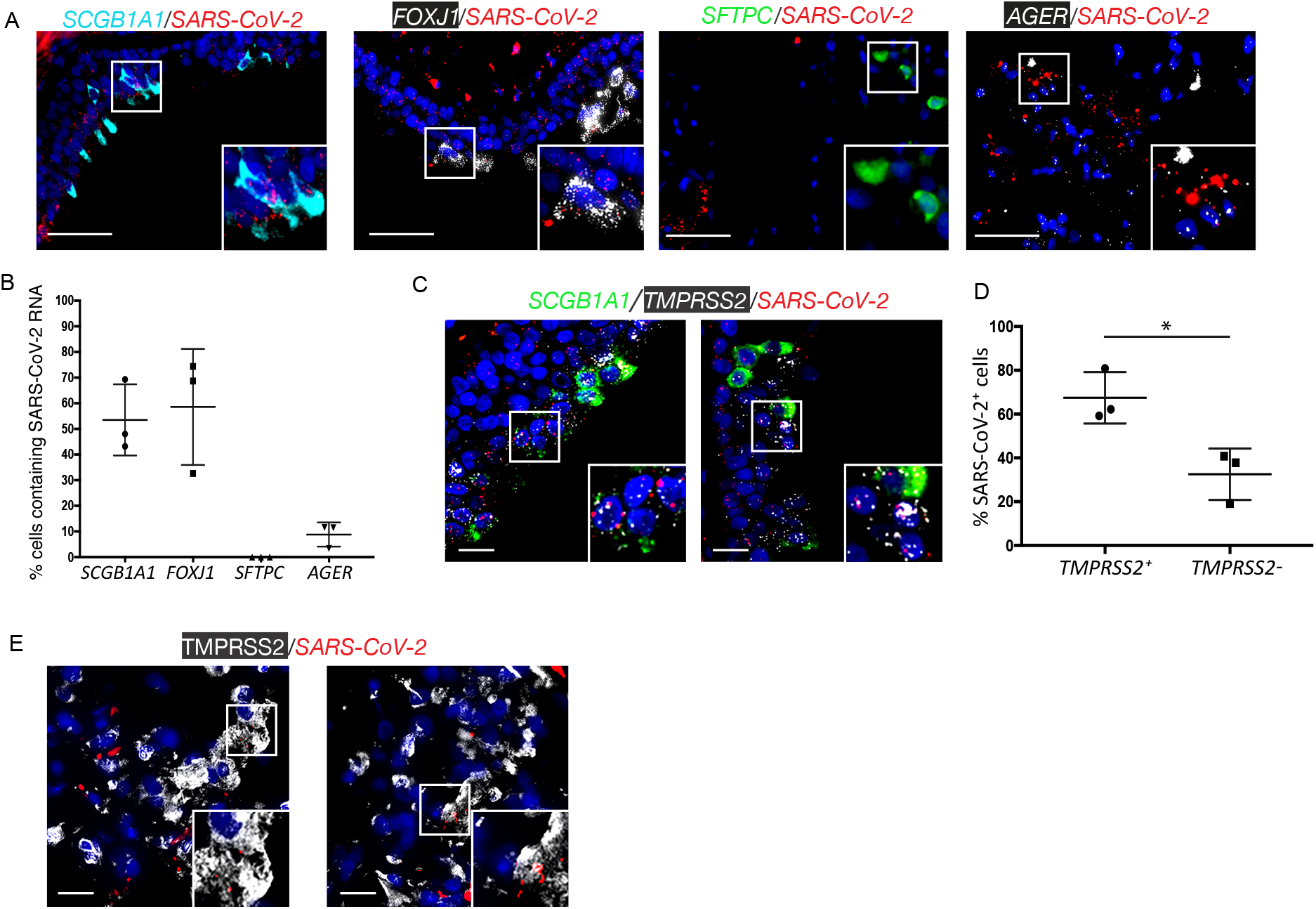
Spatial localization of SARS-CoV-2 RNA in lung autopsy tissue from fatal COVID-19. A) RNA-ISH of SARS-CoV-2 RNA (red) with epithelial cell markers *SCGB1A1 (*secretory cells, cyan), *FOXJ1* (ciliated cells, white), *SFTPC* (AT2 cells, green), *AGER* (AT1 cells, white); scale bar = 100μm. B) Quantification of cells containing SARS-CoV-2 RNA by epithelial subtype, as a percentage of total number of cells counted; a minimum of 150 cells were counted in each group. C)RNA ISH of large airway from same patient demonstrating RNA transcripts for *TMPRSS2* (white) in the same cells containing SARS-CoV-2 RNA (red), with secretory cells labeled in green (*SCGB1A1)* for context; scale bar = 100μm. D) Quantification of *TMPRSS2* expression in *SARS-CoV-2+* cells, more than 1000 cells were counted from the large airways of the 3 patients, *p<0.05*. All percentages are shown in graphs with mean +/− standard deviation. E) RNA in situ hybridization (ISH) of *SARS-CoV-2* (red) with protein immunofluorescence for TMPRSS2 protein (white), scale bar = 100μm.

The epithelial cell tropism of SARS-CoV-2 is dependent upon viral attachment and activation by cell-surface proteins, e.g. ACE2 and TMPRSS2. While ACE2 has been well studied in *in vitro* platforms of differentiated lung epithelium, our findings and prior studies (15–17, 22) suggest that ACE2 is expressed at low levels in respiratory epithelium (Figures 1, S3,5). Conversely, the rising expression of *TMPRSS2* with age in both the conducting airways and alveolar epithelium may explain the increased susceptibility to symptomatic infection and severe ARDS observed in adults relative to children. To date, there have been very few cases of vertical transmission of SARS-CoV-2 to newborns(23), despite viral particles being identified in the placentas of women who were infected with SARS-CoV-2(24). In light of the very low expression of *ACE2* in ciliated epithelial cells, our data indicate that ACE2-independent attachment may play a role in SARS-CoV-2 cellular entry and pathogenesis. CD147 (basigin, *BSG*) has also been proposed as an alternative SARS-CoV-2 receptor(25) and is expressed broadly including in ciliated cells in which its expression increased across developmental time; in contrast, Bsg expression decreased in the other epithelial cell types (Fig. S6A-B). Alternatively, recent data indicate that ACE2 may be transiently induced by interferon signaling to facilitate cellular entry, with increased interferon signaling after initial viral entry propagating ACE2 expression in neighboring cells(26). Across developmental time in the lung epithelium, we observed a progressively increasing interferon signature with increasing age, most prominent in AT2 and AT1 cells (Fig. S7A-B). This raises the possibility that the developing lung may have relatively less interferon “priming” and therefore have less efficient ACE2 induction to serve as the primary entry receptor to facilitate alveolar infectivity. In contrast, we did not observe developmental differences in basal expression of innate inflammatory genes linked to coronaviruses response including: *Irf3, Tnf, Mapk1/3, Nfkb1/2, and Cxcl15* (26–29) (Fig. S7C).

The very low levels of *TMPRSS2* expression in human infants, and nearly absent levels of expression of *Tmprss2* in prenatal mice, suggest a mechanism by which neonates may be relatively protected against severe forms of COVID-19. After the SARS outbreak in 2003, several groups(30–33) reported that proteolytic cleavage of the SARS-CoV spike (S) protein is a prerequisite for viral activation and host cell entry, with SARS-CoV tropism for TMPRSS2-expressing cells in primates(34). Prior work studying SARS-CoV has shown that inhibition of TMPRSS2 with the serine protease inhibitor Camostat, partially prevented SARS-CoV infection(35), and addition of a cathepsin inhibitor (Aloxistatin) to Camostat potentiated the antiviral effect(35).These data highlight the potential for TMPRSS2 inhibition as an effective strategy for reducing SARS-CoV-2 infection, although it is important to note this does not suggest TMPRSS2 is required for for SARS-CoV-2 entry *in vivo*.

In summary, these data suggest that developmental regulation of viral entry-factors may be the primary determinant of the age-related differences in SARS-CoV-2 susceptibility and severity. The identification of changes in *TMPRSS2* expression associated with age presents a biological rationale for the observed rarity of severe lower respiratory tract SARS-CoV-2 disease in children, and underscores the opportunity to consider *TMPRSS2* inhibition as a potential therapeutic target for SARS-CoV-2.

## METHODS

Methods are described in detail in the supplement.

### Data availability

Raw data and counts matrices used in generating this dataset are available through the Gene Expression Omnibus (GEO) (pending). Code used for dataset integration and analyses in this manuscript are available at https://github.com/KropskiLab.

### Subjects and samples

For the use of human tissue for research, all studies were approved by the Vanderbilt Institutional Review Board (VanderbiltIRB #’s 060165, 171657, 200490).

### Animal care

The protocol for this was approved by the Institutional Animal Care and Use Committee of Vanderbilt University (Nashville, TN) and was in compliance with the Public Health Services policy on humane care and use of laboratory animals.

## Acknowledgements

This work was supported by NIH K08143051 (JMS), K08HL130595 (JAK), R01HL145372(JAK/NEB), P01HL092470(TSB), K08HL127102(EJP), K08HL133484(JTB), R01AI077505 (DWH), P30AI110527 (SAM), R01AI142095(SAK/SAM), TR002243. Flow Cytometry experiments were performed in the VMC Flow Cytometry Shared Resource. The VMC Flow Cytometry Shared Resource is supported by the Vanderbilt Ingram Cancer Center (P30 CA68485) and the Vanderbilt Digestive Disease Research Center(DK058404).

## Consortia

### Vanderbilt COVID-19 Cohort Consortium (VC3)

Justin L. Balko, Suman Das, David Haas, Spyros A. Kalams, Jonathan A. Kropski, Christine Lovly, Simon A. Mallal, Elizabeth J. Phillips, Wei Zheng

### Human Cell Atlas Lung Biological Network

The members of HCA Lung Biological Network are Nicholas E. Banovich, Pascal Barbry, Alvis Brazma, Tushar Desai, Thu Elizabeth Duong, Oliver Eickelberg, Christine Falk, Michael Farzan, Ian Glass, Muzlifah Haniffa, Peter Horvath, Deborah Hung, Naftali Kaminski, Mark Krasnow, Jonathan A. Kropski, Malte Kuhnemund, Robert Lafyatis, Haeock Lee, Sylvie Leroy, Sten Linnarson, Joakim Lundeberg, Kerstin B. Meyer, Alexander Misharin, Martijn Nawijn, Marko Z. Nikolic, Jose Ordovas-Montanes, Dana Pe’er, Joseph Powell, Stephen Quake, Jay Rajagopal, Purushothama Rao Tata, Emma L. Rawlins, Aviv Regev, Paul A. Reyfman, Mauricio Rojas, Orit Rosen, Kourosh Saeb-Parsy, Christos Samakovlis, Herbert Schiller, Joachim L. Schultze, Max A. Seibold, Alex K. Shalek, Douglas Shepherd, Jason Spence, Avrum Spira, Xin Sun, Sarah Teichmann, Fabian Theis, Alexander Tsankov, Maarten van den Berge, Michael von Papen, Jeffrey Whitsett, Ramnik Xavier, Yan Xu, Laure-Emmanuelle Zaragosi, and Kun Zhang. Pascal Barbry, Alexander Misharin, Martijn Nawijn, and Jay Rajagopal serve as the coordinators.

## Author Contributions

JMS and JAK conceived the study. EJP, JB, CJT, CJ, AH, PG, DSN performed experiments. BAS, ACH, NEB, JMS, JAK analyzed transcriptomic data. MK, SHG, VC3, TSB provided samples, BAS, ACH, JAK and JMS interpreted data with assistance from TSB, SHG, SHG, SAW, NEB, LZB, MHK, VC3 and the HCA Lung Biological Network. BAS, LTB, JAK and JMS wrote the manuscript and all authors provided critical feedback on the manuscript.

## Competing Interests

JAK has received advisory board fees from Boehringer Ingelheim, Inc, and has research contracts with Genentech. TSB has received advisory board fees from Boehringer Ingelheim, Inc, Orinove, GRI Bio, Morphic, and Novelstar, and has research contracts with Genentech and Celgene.

## Supplementary Materials

Detailed Methods

Figures S1–9

## Methods

### Animal Care and Tissue Fixation

C57BL/6 mice were used for all experiments. Timed matings were performed as previously described(1) and mice were sacrificed at P0, P7, P14, or P64 for single cell RNA sequencing or lung block fixation. E18 lungs were isolated by removing pups from the mouse uterus and isolating lung tissue. E18 and P0 lungs were fixed in formalin. P7, P14, P64, 12 month, and 24 month old mouse lungs were inflation-fixed by gravity filling with 10% buffered formalin and paraffin embedded. This protocol was approved by the Institutional Animal Care and Use Committee of Vanderbilt University (Nashville, TN) and was in compliance with the Public Health Services policy on humane care and use of laboratory animals.

### Single cell isolation and flow cytometry

At the indicated timepoints, lung lobes were harvested, minced, and incubated for 30 minutes at 37°C in dissociation media (RPMI-1640 with 0.7 mg/ml collagenase XI and 30 mg/ml type IV bovine pancreatic DNase). After incubation, lobes were passed through a wide bore pipet tip and filter through a 40 mm filter. Single cell lung suspension was then counted, aliquoted, and blocked with CD-32 Fc block (BD cat #553142) for 20 minutes on ice. After 2% FBS staining buffer wash, cells were incubated with the conjugated primary antibodies anti-CD45 (BD cat # 559864) and anti-Ter119 (BD cat# 116211) as indicated below. In the same manner, fluorescence minus one controls were blocked and stained with the appropriate antibody controls. Cells from individual mice were then incubated with identifiable hashtags, resuspended in staining buffer, and treated with PI viability dye. CD45 negative, Ter119 negative, viable cells were collected by fluorescence associated cell sorting using a 70 mm nozzle on a 4-laser FACSAria III Cell Sorter. Both single and fluorescence-minus-one controls were used for compensation.

### scRNA-seq library preparation and next-generation sequencing

ScRNA-seq libraries were generated using the 10X Chromium platform 5’ library preparation kits (10X Genomics) following the manufacturer’s recommendations and targeting 10,000 - 20,000 cells per sample. Next generation sequencing was performed on an Illumina Novaseq 6000. Reads with read quality less than 30 were filtered out and CellRanger Count v3.1 (10X Genomics) was used to align reads onto mm10 reference genome.

### Ambient RNA filtering

Ambient background RNA were cleaned from the scRNA-seq data with “SoupX” (version 1.2.2, Wellcome Sanger Institute, Hinxton, Cambridgeshire, UK); (2)) in RStudio (version 1.2.5001, RStudio, Inc., Boston, Massachusetts, USA). Matrix files from CellRanger were read into the global environment and combined into a “SoupChannel.” Data from the SoupChannel were passed to Seurat (version 3.1.4, New York Genome Center, New York City, New York, USA)(3, 4) for data normalization, identification of variable features, data scaling, principal component analysis, uniform manifold approximation and projection (UMAP) for dimensionality reduction, calculation of nearest neighbors, and cluster identification. The SoupX pipeline was used for each time point to determine which genes were most likely contributing to the ambient background RNA. We utilized the following genes to estimate the non-expressing cells, calculate the contamination fraction, and adjust the gene expression counts: *Dcn, Bgn, Aspn, Ecm2, Fos, Hbb-bs, Hbb-bt, Hba-a1, Hba-a2, Lyz1, Lyz2, Mgp, Postn, Scgb1a1*. For time points that were sorted to select for epithelial cells, the following genes were added to the SoupX pipeline: *Sftpc, Hopx, Ager, Krt19, Cldn4, Foxj1, Krt5, Sfn, Pecam1*. New matrix files were created by SoupX and used for subsequent analyses in Scanpy.

### Data integration and clustering

Data integration and clustering was performed using a standard Scanpy workflow (Scanpy v1.46)(5). Individual SoupX “cleaned” libraries were concatenated and analyzed jointly. After quality filtering (removal of cells with <500 or >5000 genes, or >10% mitochondrial gene expression), data were normalized, log-transformed, highly-variable genes were identified, and percent mitochondrial gene expression and and read depth were regressed. Following data scaling, and principal components analysis, Leiden clustering was performed followed by visualization of canonical marker genes. Red blood cells and doublet clusters (containing non-physiologic marker combinations, i.e. *Epcam*+/*Pecam1*+) were filtered. Batch correction for dataset integration was performed using batch-balanced K-nearest neighbors(6). Principal components 1:30 were used for clustering and Uniform Manifold Approximation and Projection (UMAP) embedding(7). Leiden clustering (resolution 0.8) was then performed on the integrated dataset and cell-types assigned based on marker expression profiles.

### Subjects and samples

Lung tissue from 20 healthy human subjects was obtained at the time of surgical biopsy or autopsy, with death occurring from non-respiratory causes. Subjects were classified as infants (between birth and age 2 years), children (3-17 years), and adult (54-69 years old). Adult tissue included male and female donors, and all were lifetime nonsmokers. SARS-CoV-2 lung tissue was obtained at the time of autopsy from 3 patients who died from hypoxic respiratory failure (ages 51, 82, 97). All studies were approved by the Vanderbilt Institutional Review Board (VanderbiltIRB #’s 060165, 171657, 200490)

### RNA *in situ* hybridization

RNAScope technology (ACDBio) was used to perform all RNA *in situ* hybridization (RNA ISH) experiments according to manufacturer’s instructions. RNAScope probes to the following human genes were used: *TMPRSS2, ACE2, SCGB1A1, FOXJ1, SFTPC,* and *AGER*. Probes to the following mouse genes were used: *Tmprss2, Ace2, Scgb1a1, Foxj1, Sftpc, Hopx.* RNAScope probe to the SARS-CoV-2 virus (sense) was also used. Our validation of the SARS-CoV-2 probe included application of this probe to lung tissue from healthy adult lungs from a patient who died in 2018, which showed no amplification or fluorescence (Figure S8). Positive control probe (*PPIB)* and negative control probe (*DapB)* were purchased from the company and performed with each experimental replicate, with representative data shown (Figure S8).

### Immunofluorescence

Using the published protocol from ACD Bio (https://acdbio.com/technical-support/user-manuals, 323100-TN), we combined immunofluorescence with RNA in situ hybridization. We used a monoclonal antibody to TMRPSS2 (Abcam, ab109131) and RNAScope probes to *TMPRSS2 and FOXJ1* in human samples and *Tmprss2* and *Foxj1* in murine samples. Nuclei were stained with DAPI (Vector Laboratories).

### Image acquisition and analysis

Fluorescent images were acquired using a Keyence BZ-X710 with BZ-X Viewer software with 40X objective. Wavelengths used for excitation included 405, 488, 561, and 647nm lines. Automated image analysis was performed with Halo software (Indica Labs); an example of cellular segmentation and determination of co-localization is demonstrated in Figure S9. Cell area of *Tmprss2/TMPRSS2* probe was calculated as a percentage of total cell area for each epithelial subtype. Number of *Tmprss2/TMPRSS2* positive cells (defined as having 5 or more copies of *Tmprss2/TMPRSS2* by Halo analysis) as a percentage of total epithelial cells was also calculated. Presence of SARS-CoV-2 in epithelial cells was determined by Halo segmentation and analysis.

### Statistical approach

Statistical tests used for specific comparisons are described in the relevant figure legends. Comparisons of gene expression from scRNA-seq were performed by analysis of variance (anova) using the linear model (lm) function in R version 3.6.3.

## Supplementary Figures

**Figure S1.**
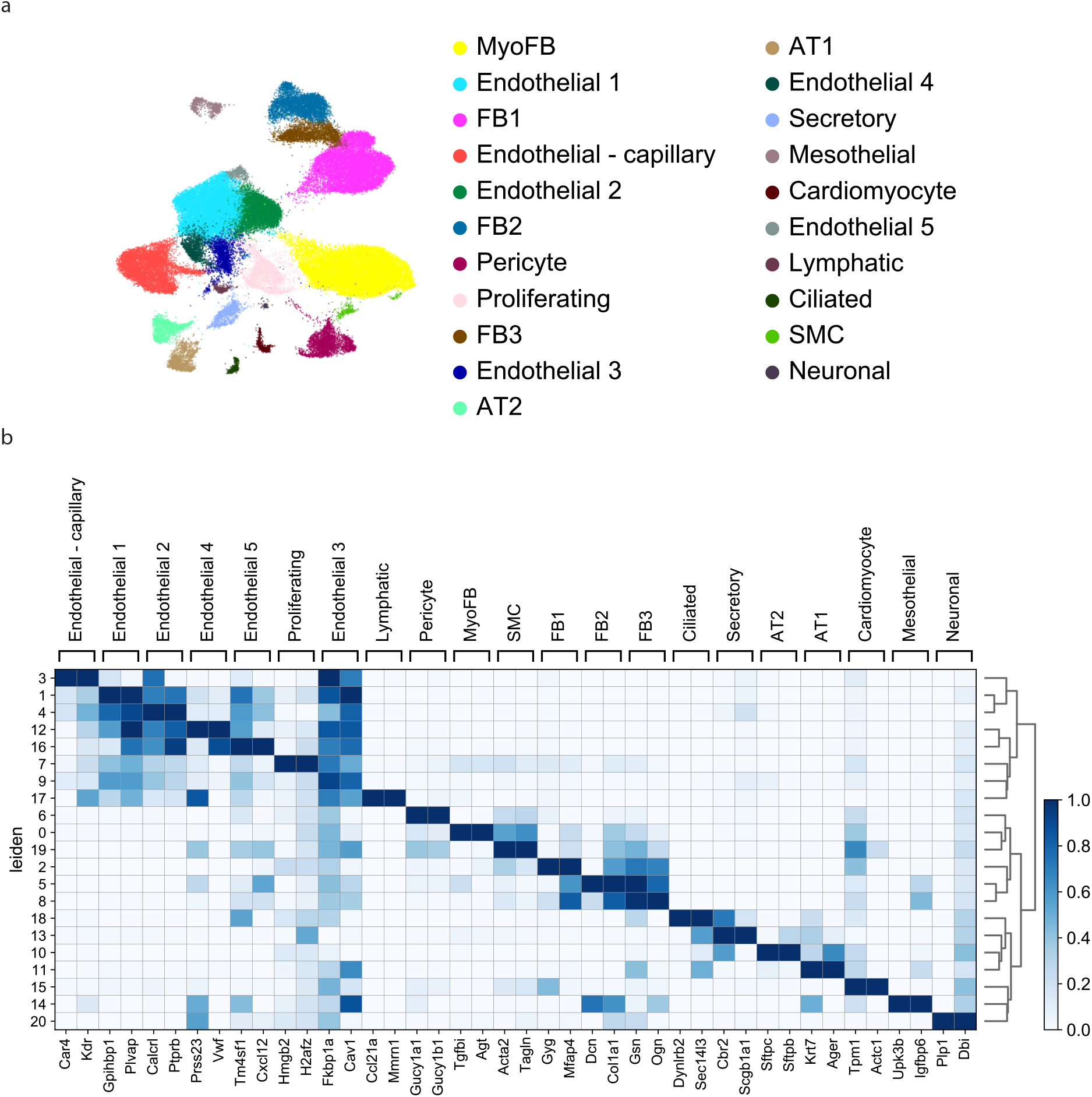
Cell types identified by time-series scRNA-seq. A) Enlarged UMAP embedded of 67,629 cells from across developmental time annotated by cell-type. B) Matrixplot depicting the top 2 markers for each cell-type determined by Wilcoxon-test comparing a given cluster to all others.

**Figure S2:**
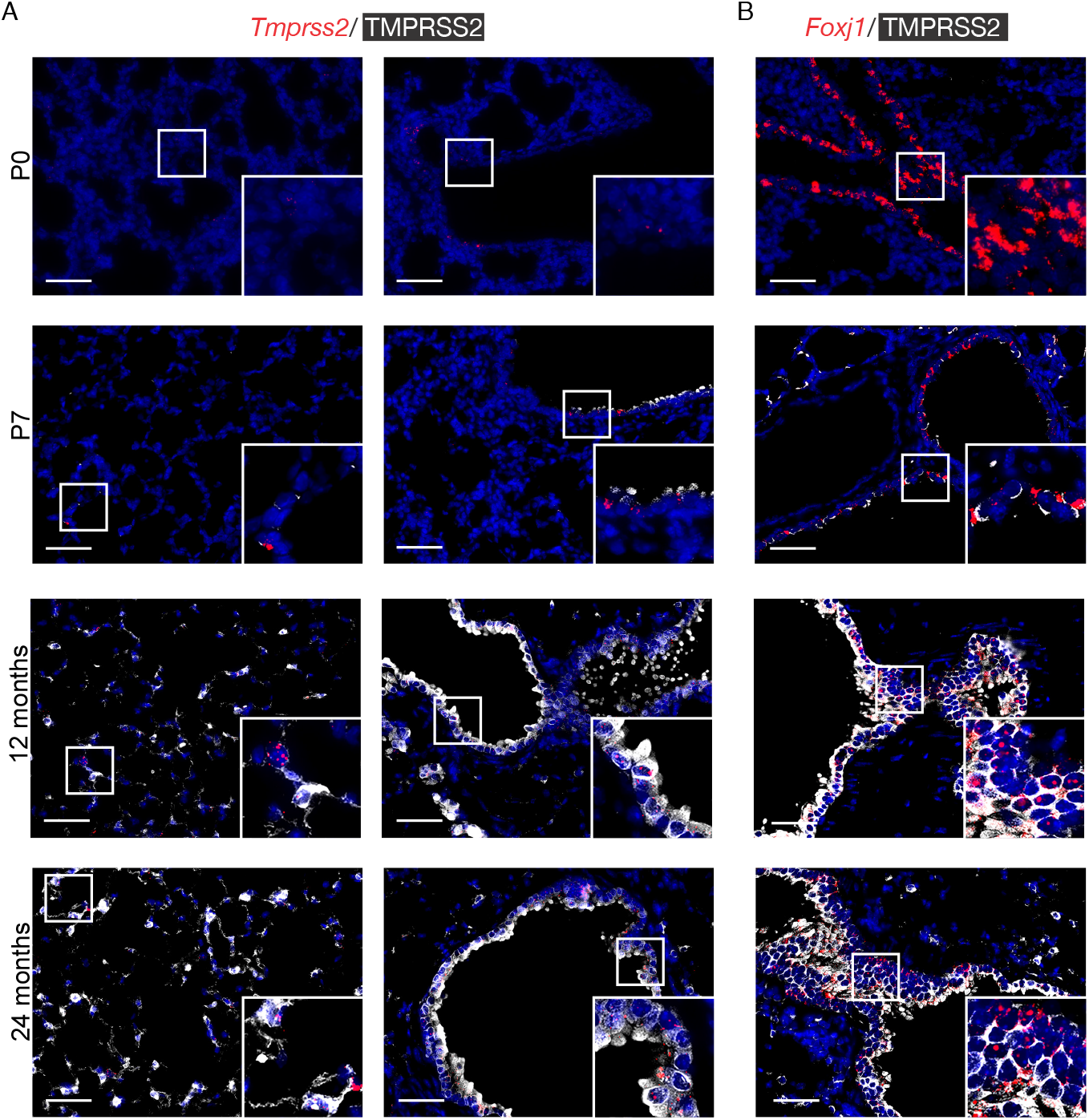
Spatial and temporal localization of TMPRSS2 across murine lung development. A) RNA in situ hybridization (ISH) of *Tmprss2* expression (red) with protein immunofluorescence for TMPRSS2 protein (white). B) RNA ISH of *Foxj1* expression (red) with protein immunofluorescence for TMPRSS2 protein (white). Formalin fixed paraffin embedded tissue from lungs at timepoints P0, P7, 12 months, 24 months. Lungs from 3 mice at each time point were used, with ten 40X images obtained per slide. Scale bar = 100um.

**Figure S3:**
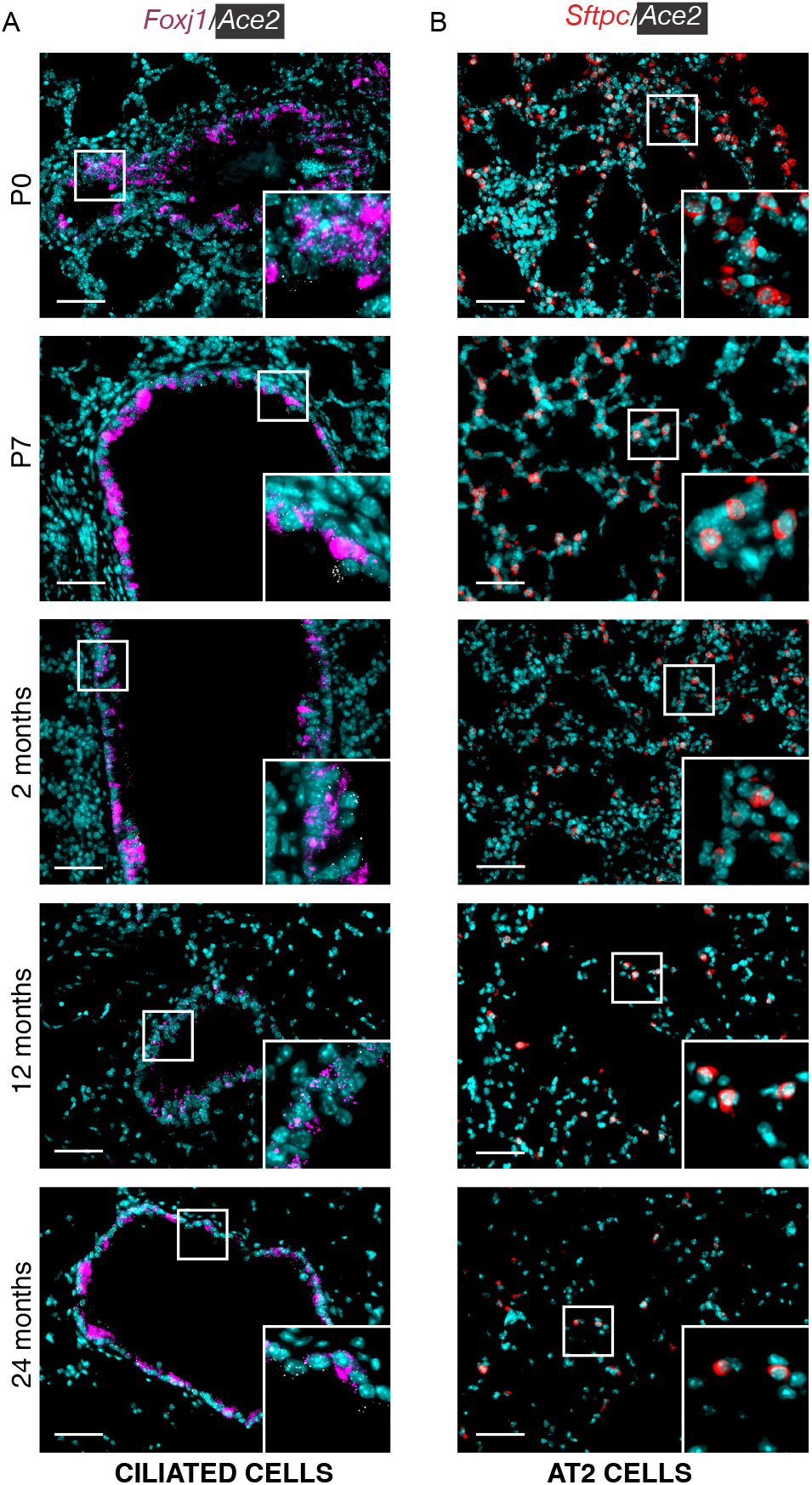
Spatial and temporal localization of *Ace2* expression across lung development. A) RNA in situ hybridization (ISH) of *Ace2* expression (white) with epithelial cell markers *Foxj1* (ciliated cells, magenta) and *Sftpc* (surfactant protein C, AT2 cells, red). Formalin fixed paraffin embedded tissue from lungs at timepoints P0, P3, P5, P7, P14, P64. Lungs from 3 mice at each time point were used, with five 40X images obtained per slide. Scale bar = 100um.

**Figure S4:**
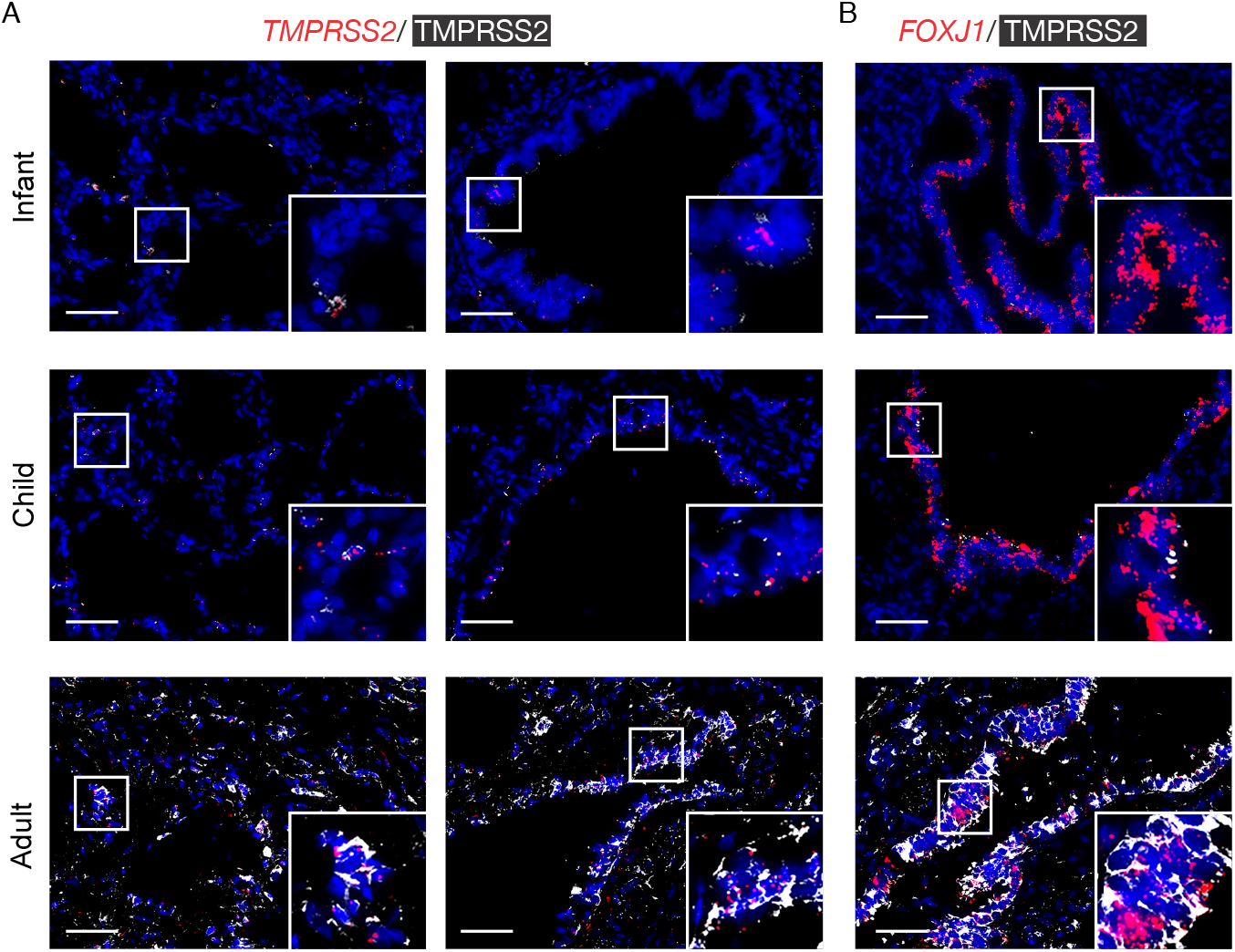
Spatial and temporal localization of TMPRSS2 across the human lifespan. A) RNA in situ hybridization (ISH) of *Tmprss2* expression (red) with protein immunofluorescence for TMPRSS2 protein (white). B) RNA ISH of *Foxj1* expression (red) with protein immunofluorescence for TMPRSS2 protein (white). Formalin fixed paraffin embedded tissue from lungs between the ages of birth and 69 years was analyzed. We defined infants as birth-2 years, children 3years-17 years, and adult specimens were between 53-69 years of age. Five 40X images obtained per slide for analysis. Scale bar = 100 μm.

**Figure S5:**
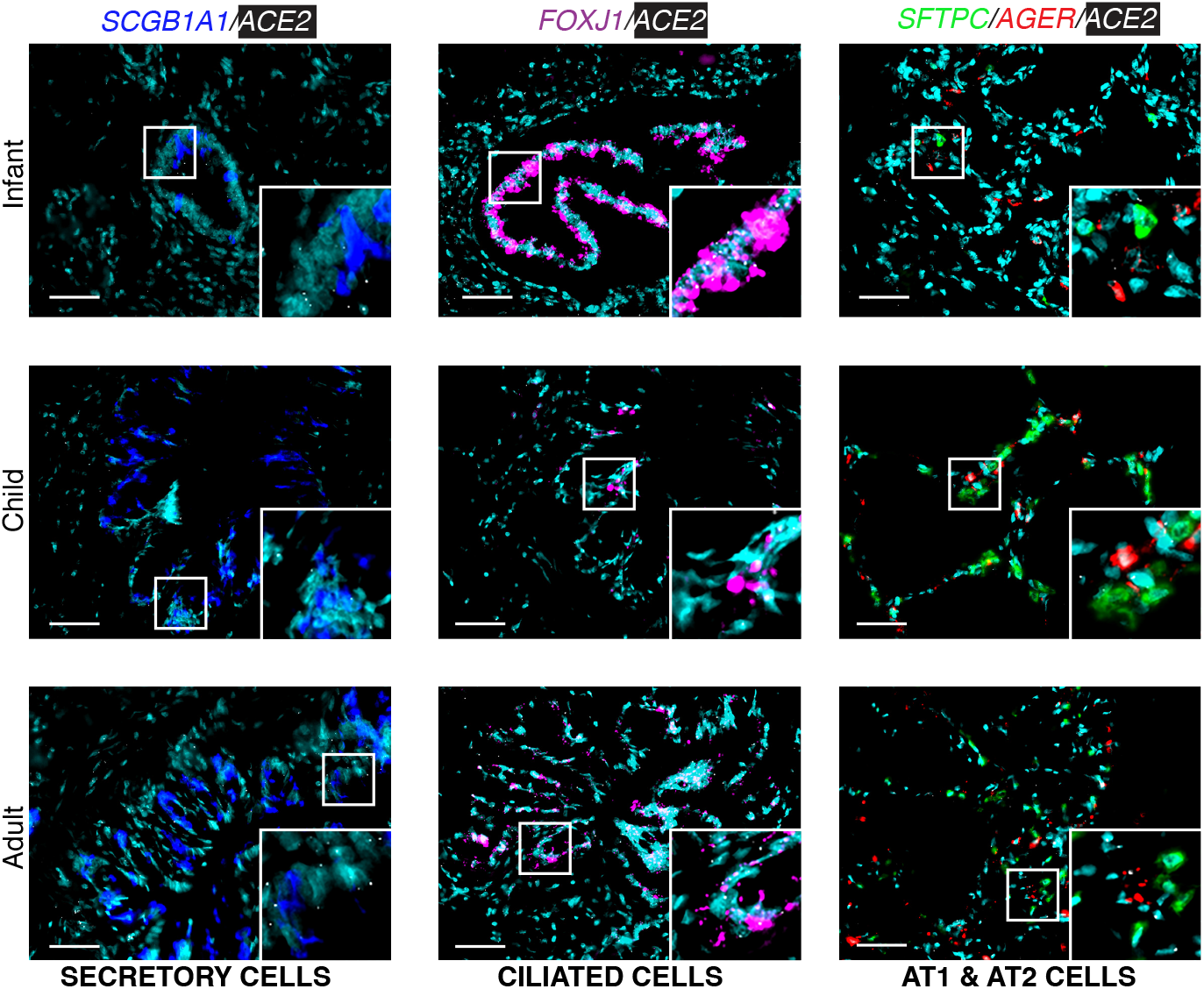
Spatial and temporal localization of *ACE2* expression in the lung across the human lifespan. A) RNA *in situ* hybridization (ISH) of *ACE2* expression (white) with epithelial cell markers *SCGB1A1* (secretory cells, blue), *FOXJ1* (ciliated cells, magenta), *SFTPC* (surfactant protein C, AT2 cells, green), *AGER* (AT1 cells, red). Formalin fixed paraffin embedded tissue from 25 human lungs between the ages of birth and 69 years was analyzed. We defined infants as birth-2 years, children 3years-17 years, and adult specimens were between 53-69 years of age. Five 40X images obtained per slide for analysis. Scale bar = 100 μm.

**Figure S6.**
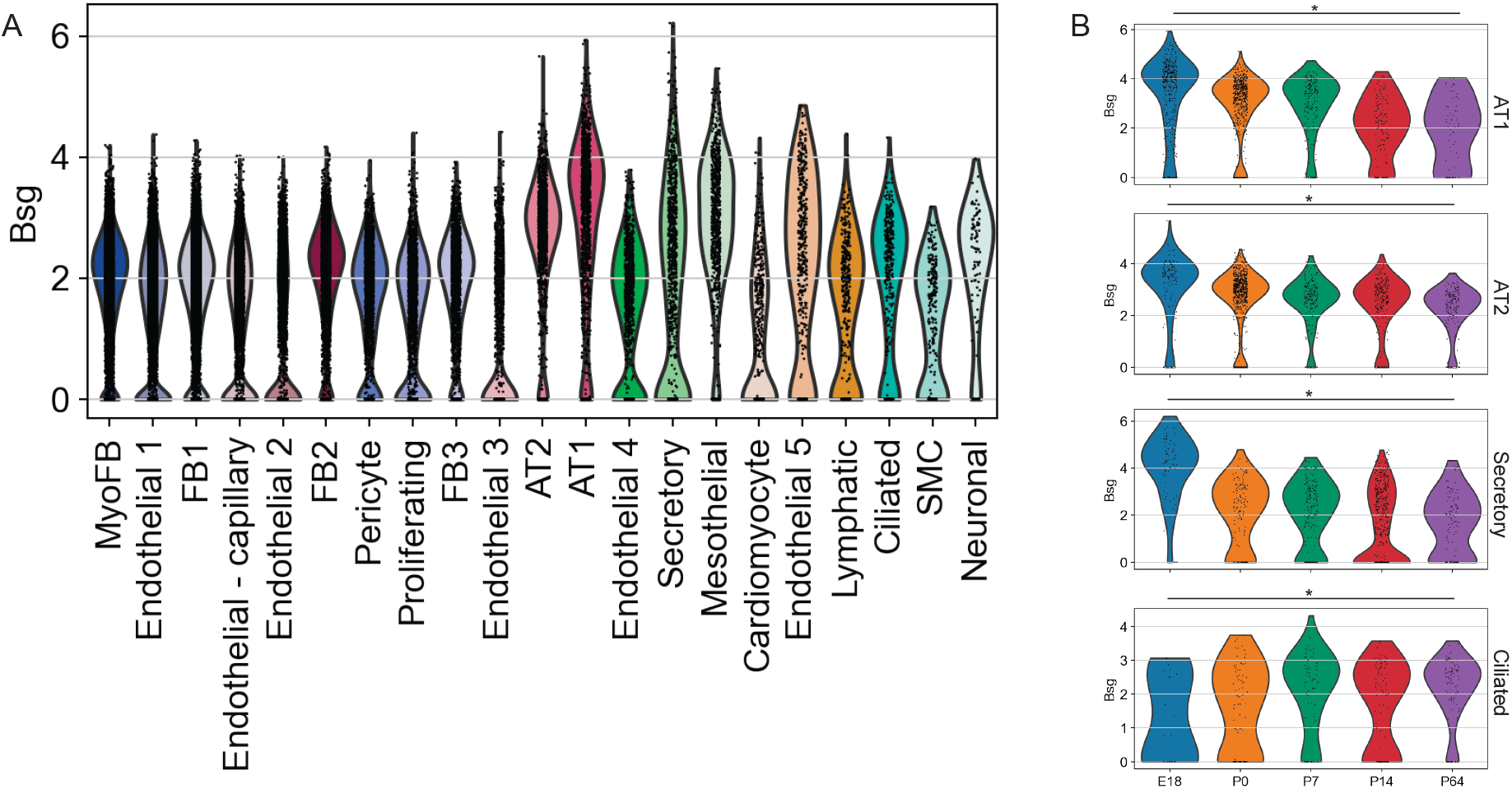
Expression of proposed alternative SARS-CoV-2 receptor CD147 (Bsg) across lung development. A) Violin plot depicting *Bsg* expression across cell types. B) Violin plots depicting *Bsg* expression in epithelial cell types across developmental time. * p <0.05 by ANOVA with Bonferroni correction for multiple testing.

**Figure S7.**
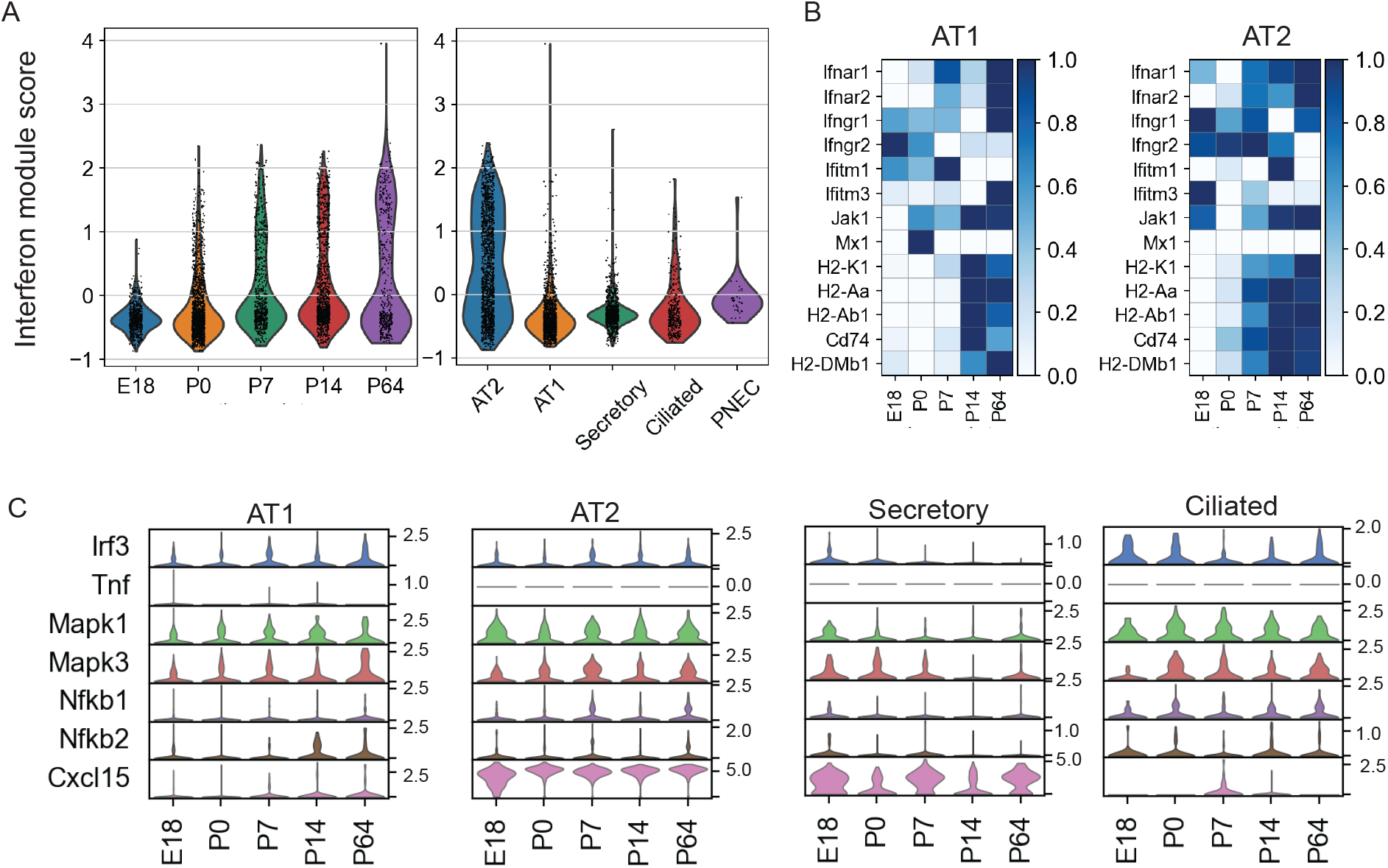
Immune expression programs across developmental time. Interferon gene module score was calculated using the scoregenes tool in Scanpy v1.46 using interferon receptors and canonical interferon-response genes plotted A) in all epithelial cells by developmental timepoint and B) by epithelial cell type. C) Heatmap depicting relative expression of interferon-related genes in AT2 and AT1 cells across developmental time. D) Expression of innate immune gene across developmental time.

**Figure S8:**
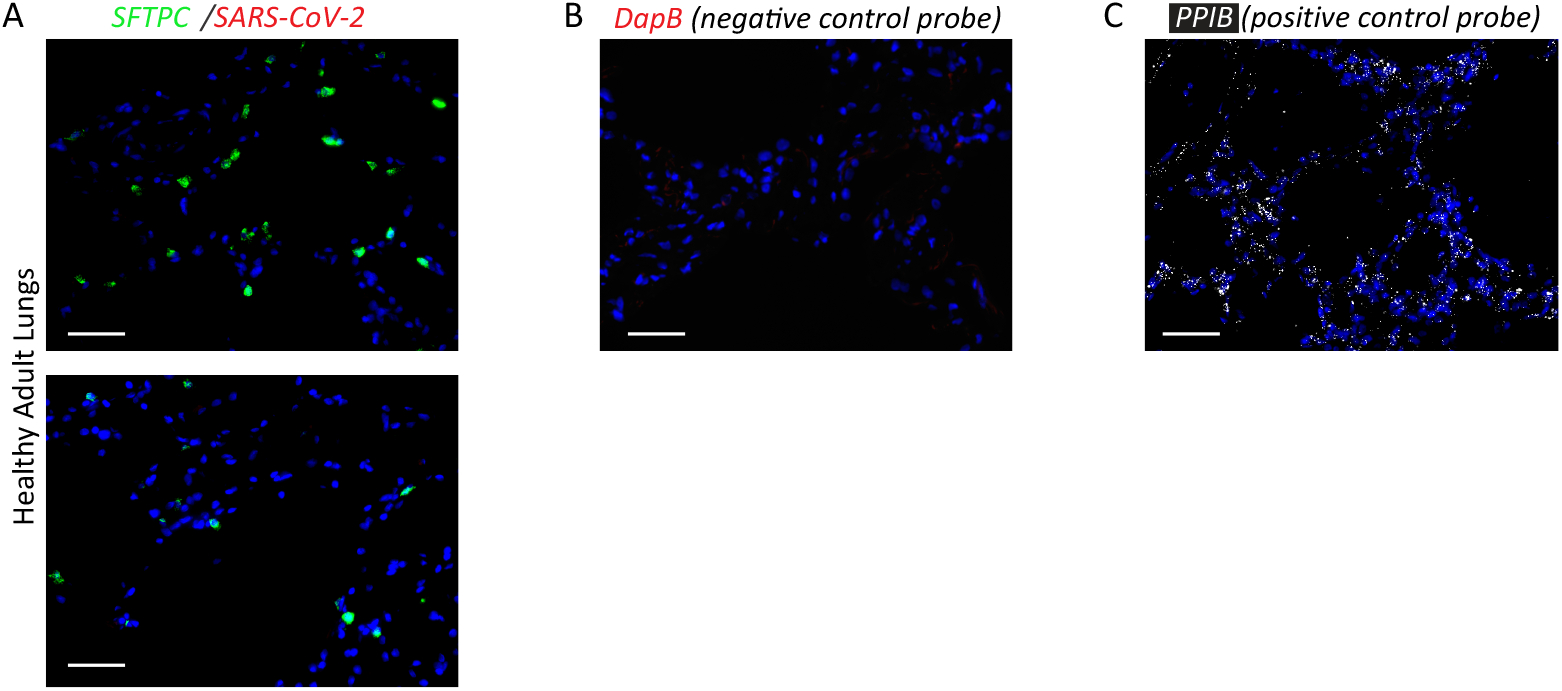
A) RNA *in situ* hybridization (ISH)with probes for SARS-CoV-2 RNA (red) and *SFTPC* (green) in lungs from a patient who died in 2018, which demonstrates no amplification or fluorescence of SARS-CoV-2 RNA. B) RNA ISH with *DapB* (red), a negative control probe for an E.coli gene provided by ACDBio. C) RNA ISH with *PPIB* (white), a positive control probe for a housekeeping gene provided by ACDBio. Scale bar = 100 μm.

**Figure S9:**
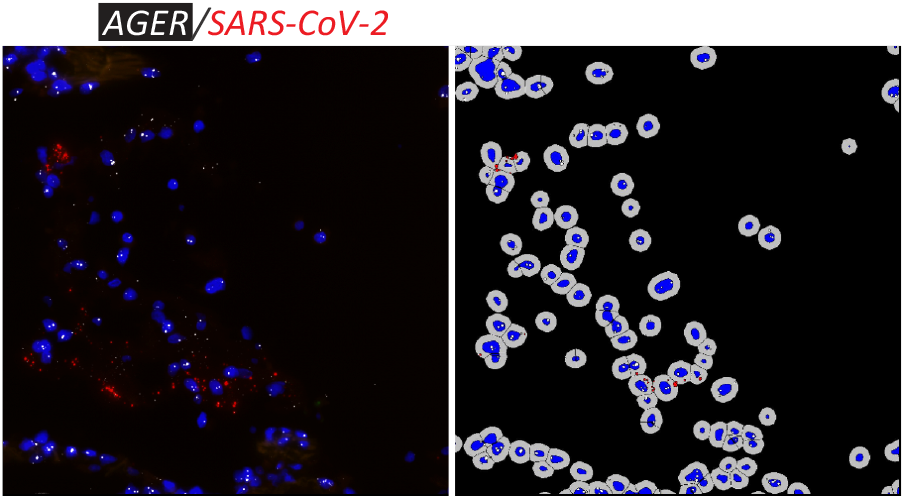
Example of cellular segmentation and automated labeling of epithelial cells as containing SARS-CoV-2 virus, shown here with *AGER* positive cells from lung parenchyma of a patient who died of COVID-19.

## Notes

### Summary of Updates

Revised version of the manuscript includes protein-level data and quantification of SARS-CoV-2 virus in autopsy tissue from n=3 samples.

